# Label-Free Nucleoli Measurement by 3D Holo-Tomographic Flow Cytometry Using Biolens Phase Compensation

**DOI:** 10.64898/2026.01.12.699016

**Authors:** Daniele Pirone, Michela Schiavo, Giusy Giugliano, Sandro Montefusco, Annalaura Montella, Martina Mugnano, Vincenza Cerbone, Maddalena Raia, Giulia Scalia, Vittorio Bianco, Lisa Miccio, Mario Capasso, Achille Iolascon, Diego Luis Medina, Pasquale Memmolo, Demetri Psaltis, Pietro Ferraro

## Abstract

Holo-Tomographic Flow Cytometry (HTFC) holds the potential to transform cellular research and clinical screening through 3D label-free quantitative phase imaging (QPI) of flowing single cells. However, it has been limited by insufficient intracellular specificity in 3D refractive index (RI) distributions, since suspended cells act as highly aberrating spherical biolenses obscuring internal structures. Here, we show the Biolens Phase Compensation (BPC), a method that corrects phase aberrations in 2D QPI projections to transform the 3D RI tomogram. Working within this new 3D pseudo-RI space demonstrates for the first time the extraction of nucleoli in HTFC. Extensive validation against 2D fluorescence flow cytometry and 3D confocal microscopy demonstrates that BPC achieves reliable intranuclear specificity. Using statistically significant single-cell analysis, we provide multiplexed quantitative 3D measurements of nested intracellular compartments (cytoplasm, nucleoplasm, nucleoli). This approach extends label-free HTFC toward capabilities of gold-standard Fluorescence Microscopy, overcoming its well-known drawbacks in intracellular and intranuclear staining.

## Introduction

To comprehend the complexities of cell biology in health and disease, researchers utilize a variety of imaging techniques [1], each with distinct advantages and limitations. Among them, Fluorescence Microscopy (FM) enables real-time visualization of specific intracellular components, yet its resolution is often insufficient for detailed structural analysis. Moreover, using fluorescently labeled antibodies or dyes can lead to photobleaching, phototoxicity, or altered cell behavior, affecting organelle detection and introducing artifacts [2,3]. The field therefore still faces challenges in developing non-invasive, label-free imaging methods that provide comprehensive insights into intracellular structures and functions without perturbing cellular physiology.

To address these limitations, label-free optical imaging techniques are rapidly gaining ground [4]. By eschewing chemical stains and instead exploiting inherent variations in refractive index (RI) as endogenous markers, Quantitative Phase Imaging (QPI) offers a compelling alternative [5]. Among QPI approaches, Holographic Tomography (HT) is particularly powerful and promising, probing samples from multiple angles to reconstruct the volumetric RI structure [6]. The recent development of Holo-Tomographic Flow Cytometry (HTFC) has enabled 3D quantitative imaging of flowing single cells [7-11], providing detailed morphological and biophysical measurements without fluorescent labels. However, despite advances, QPI has not achieved the intracellular specificity of established FM. Although 3D RI distributions contain information on intracellular structures, mapping them directly to specific organelles is challenging in the absence of exogenous markers. Significant efforts have addressed this gap [12], mainly using deep learning to introduce “virtual specificity” [13], which has also been extended to 3D RI tomograms of adherent cells in HT [14,15]. However, in microfluidic environments such as HTFC, training deep convolutional neural networks to map organelle RIs to fluorescence signals is cumbersome, due to the complexity of acquiring co-registered multimodal datasets of flowing cells. To circumvent this obstacle, recently we have introduced the Computational Segmentation based on Statistical Inference (CSSI) method for identifying intracellular organelles within HTFC reconstructions, avoiding both virtual staining and the complexities of multimodal imaging acquisition [16]. CSSI employs a robust clustering algorithm to detect statistical similarities among groups of intracellular RI voxels. It was first demonstrated for segmenting convex nuclei in mammalian cells [16], later extended to reshaped vacuoles in yeast [17] and concave cup-like nuclei associated with genotypic mutations in leukemia [18]. Further applications include detection and isolation of lysosomes using CSSI-based nucleus knowledge [19]. However, achieving full specificity in HTFC remains a major challenge. The diversity of subcellular structures extends beyond those currently measurable, and existing label-free approaches are insufficient to resolve complex organelles such as nucleoli. The nucleolus, a distinct nuclear organelle, is primarily responsible for ribosomal RNA synthesis and ribosome subunit assembly [20]. Beyond this canonical role, it contributes to nuclear organization, cell-cycle regulation, and the cellular stress response [20,21]. Changes in nucleolar size, number, or morphology have been associated with pathological states, including cancer, neurodegenerative disorders, and viral infections [22]. In particular, nucleolar enlargement correlates with increased proliferation in cancer, whereas reduced nucleolar size is linked to aging and diseases such as Alzheimer’s and Parkinson’s [23-25].

In this work, we achieve intranuclear specificity in HTFC by defining a novel 3D pseudo-tomographic space, derived from the conventional 3D RI space through the proposed Biolens Phase Compensation (BPC) method illustrated in Fig. 1. Cells flowing in cytometric systems typically exhibit quasi-spherical morphologies, effectively behaving as spherical lenses [9,26]. As depicted in Fig. 1(a), this biolens effect introduces phase aberrations [27,28] in the 2D Quantitative Phase Maps (QPMs), which significantly distort the identification of intracellular compartments. Using BPC, we correct these aberrations in each single-cell 2D QPM (Fig. 1(b)). Hence, for the first time, the biolens concept is exploited within HTFC to enable accurate intracellular segmentation, tackling the particularly challenging case of nucleoli. As shown in Fig. 2, the compensated 2D QPMs are combined to reconstruct a 3D pseudo-RI tomogram, where nucleoli can be reliably localized and segmented by a BPC-based CSSI algorithm. The resulting voxel clusters are then mapped back onto the original 3D RI tomogram, enabling quantitative measurements and RI-based biophysical characterization of intracellular organelles.

**Fig. 1.**
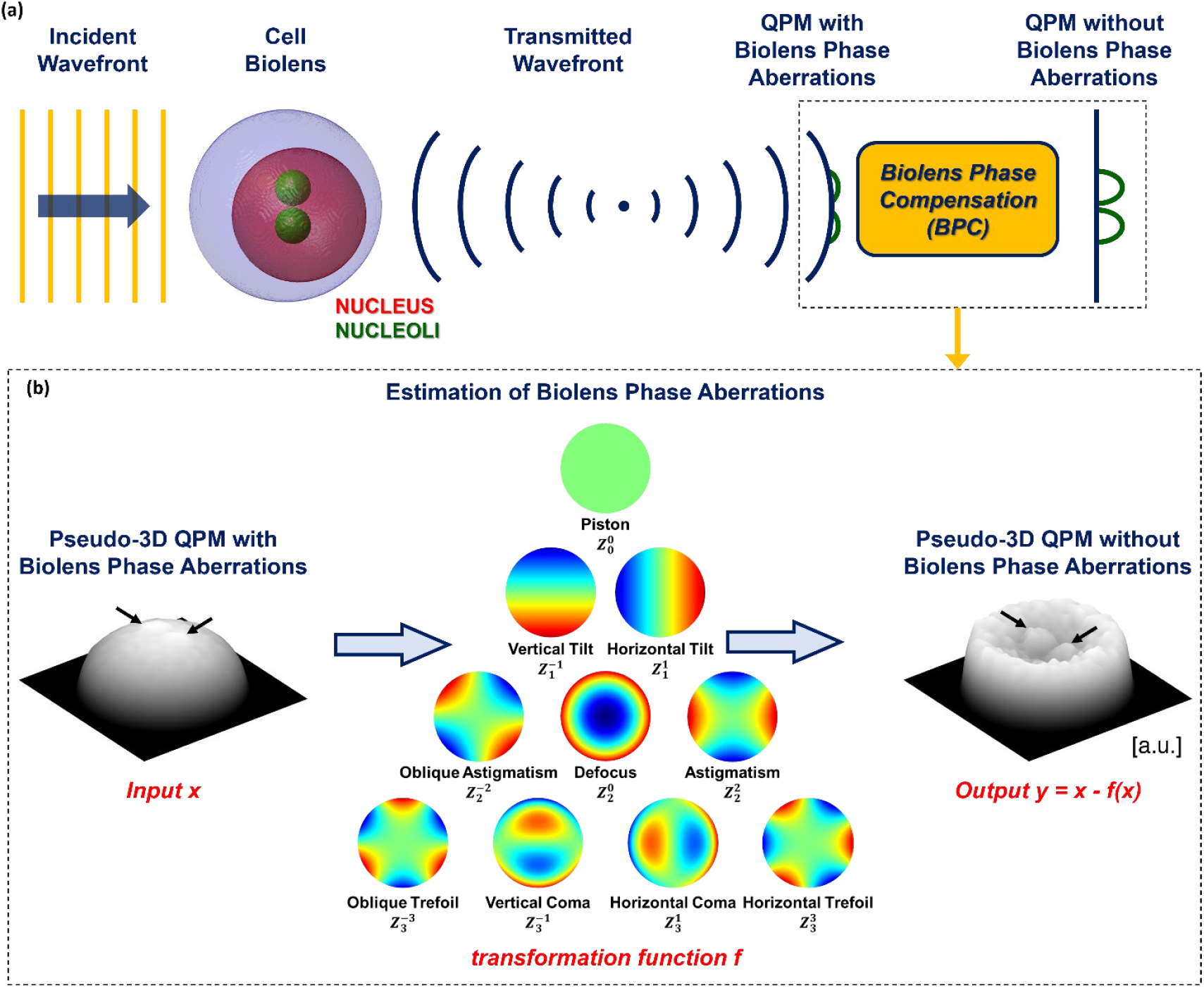
Compensation of biolens phase aberrations in single 2D QPMs. **(a)** The incident wavefront is modulated by the cell biolens into a transmitted wavefront, which is recorded as a 2D QPM. Owing to the quasi-spherical morphology of the cell biolens, phase aberrations (blue curvature) are introduced that obscure small intracellular organelles, such as nucleoli (green signals). The BPC method compensates for these biolens phase aberrations and enhances the contrast of intracellular organelles (green signals). **(b)** Sketch of the BPC method, which subtracts from each 2D QPM the biolens phase aberrations estimated directly from the same QPM, thereby revealing otherwise hidden intracellular organelles such as nucleoli (black arrows).

**Fig. 2.**
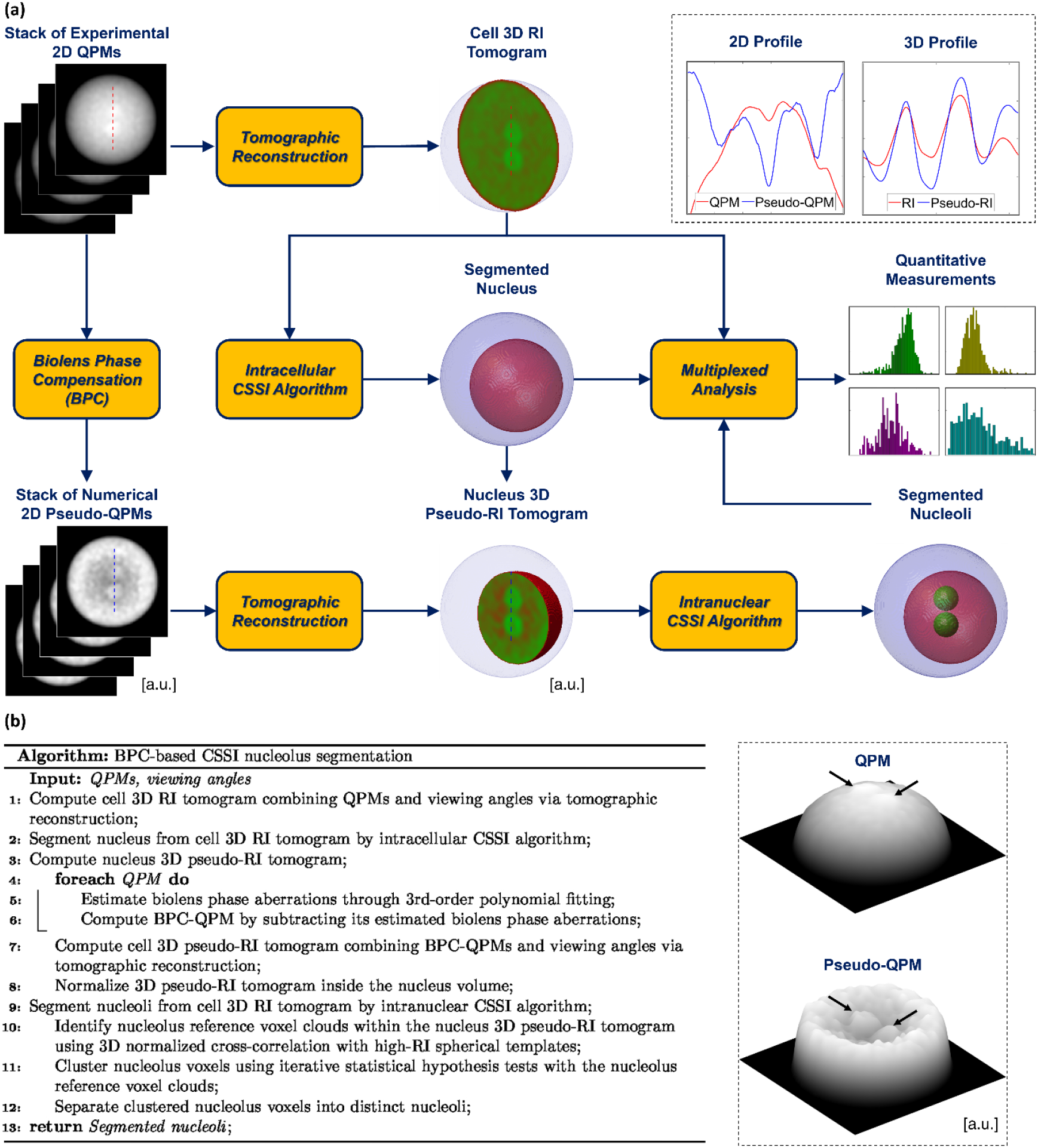
Segmentation of nucleoli in 3D RI tomograms of single cells in suspension. **(a)** Workflow of the algorithm for the nucleoli segmentation and downstream multiplexed quantitative analysis at both the intracellular and intranuclear levels. In the inset, 2D profiles (on the left) highlighted in the QPM and pseudo-QPM and 3D profiles (on the right) highlighted in the RI slice and pseudo-RI slice are compared after normalization. **(b)** Pseudo-code of the workflow in (a) for the nucleoli segmentation based on the combination between the BPC and CSSI methods. In the inset, pseudo-3D rendering of the QPM and pseudo-QPM after the subtraction of estimated biolens phase aberrations to make visible nucleoli (black arrows). In (a,b), a numerical cell phantom with two nucleoli is shown as an example.

To evaluate the method, we analyze a dataset of hundreds of HeLa cells, obtaining results consistent with literature [29]. To further validate nucleoli identification, state-of-the-art FM measurements were performed, including 2D imaging flow cytometry and 3D high-content confocal microscopy. Comparative analysis of nucleoli number and size confirm that our unstained tomographic approach performs comparably to established fluorescence-based methods, demonstrating reliable and robust intranuclear specificity. We further show that a single-cell characterization can be achieved in a multiplexed way by simultaneously measuring multiple compartments within the same cell (cytoplasm, nucleoplasm, and nucleoli) and quantifying relative morphological and biophysical features among them.

Altogether, these results indicate that an optics-based transformation of the conventional 3D RI space provides a powerful tool for investigating challenging intranuclear features in suspended cells such as nucleoli. The method advances HTFC toward full intracellular specificity by overcoming limitations inherent to multiplexed FM analyses. Efficient fluorescent staining of nucleoli typically requires nuclear membrane permeabilization, which can disrupt nuclear integrity, alter cell physiology, and increase the risk of artifacts. Thus, multiplexed labeling of nucleoplasm and nucleoli further complicates sample preparation and may bias quantitative measurements. In contrast, label-free HTFC combined with the proposed BPC-based method enables high-content multiplexed analysis of intracellular and intranuclear compartments without facing these drawbacks.

## Results and Discussion

### Nucleoli segmentation in 3D RI tomograms of label-free suspended cells

The proposed method for segmenting nucleoli in 3D RI tomograms of suspended cells is outlined in Fig. 2(a) and detailed in the pseudo-code in Fig. 2(b), using a numerical cell phantom with two nucleoli as example. In HTFC, hundreds of 2D QPMs acquired from each flowing/rotating cell are combined with their viewing angles to reconstruct the single-cell 3D RI tomogram [11]. Existing CSSI algorithms recover the 3D shape of intracellular structures occupying a single connected volume, such as the nucleus [16,18] or reshaped vacuole [17], and are therefore inadequate for organelles composed of multiple disconnected sub-volumes sharing similar RI distributions, as in the case of nucleoli [29]. Moreover, nucleoli do not occupy fixed positions within the nucleus, further complicating their localization. In fact, to implement the CSSI working principle, which relies on identifying statistical similarities within the 3D RI tomogram, a reference voxel cloud must be defined as a representative subset of the organelle volume to be segmented. Here, CSSI is first applied to isolate the nucleus, which constrains the search region for nucleolar reference voxel clouds. Although nucleoli could in principle be detected from their slightly higher RI relative to the surrounding nucleoplasm [30], the intranuclear RI contrast in HTFC is typically too low for robust identification. This limitation arises because RI reconstructions of flowing cells generally have reduced contrast compared to adherent cells under static conditions, where conventional image processing techniques can reliably detect nucleolar compartments within the thin nuclear layer [31-33]. To overcome this contrast limitation, we employ the proposed BPC method (Fig. 1) to transform RI values into a 3D pseudo-RI space, enabling accurate localization of multiple nucleolar reference voxel clouds and extending the CSSI framework to complex intranuclear structures.

According to the ray-optics approximation, each 2D QPM represents the integral of the 3D RI distribution along the optical axis. The quasi-spherical morphology of a suspended cell therefore acts as a quasi-spherical biolens, generating a quasi-paraboloid phase profile in the 2D QPM [9,26], as sketched in Fig. 1(a). This curvature, clearly visible in the pseudo-3D rendering in Fig. 1(b), obscures fine intranuclear structures such as nucleoli. To compensate for this effect, biolens phase aberrations are estimated by fitting a third-order polynomial surface (corresponding to the first three orders of Zernike polynomial terms [34]) to the recorded QPM and subtracting it from the data, yielding the compensated 2D pseudo-QPM (Fig. 1(b)). This approach is conceptually inspired by aberration compensation techniques in QPI [27,28], where system-induced aberrations are removed to improve single-cell visualization, with the key difference that here the optical system corresponds to the cell itself. Eliminating the quasi-paraboloid term noticeably enhances small intracellular features, as highlighted by the arrows in Fig. 1(b). This contrast improvement is further confirmed in the inset of Fig. 2(a), where the normalized intensity profiles of the original QPM and compensated pseudo-QPM show the enhancement provided by the BPC method.

Following the scheme in Fig. 2(a), the quasi-paraboloid subtraction is applied to all 2D QPMs of the same cell, mapping the 3D RI distribution into a new metric space consisting of a 3D pseudo-RI tomogram of the nucleus previously segmented in the 3D RI tomogram. When comparing the central slices of the RI and pseudo-RI tomograms, nucleolar compartments become markedly more distinguishable, as also shown by the normalized profiles in the inset of Fig. 2(a). This enhanced intranuclear contrast enables automatic identification of nucleolus reference voxel clouds. Nucleoli can be efficiently localized by exploiting their known properties: they are quasi-spherical [35] and exhibit the highest RIs within the nucleus [30]. Reference voxel clouds are therefore detected in the 3D BPC space using 3D normalized cross-correlation (NCC) [36] with high-RI spherical templates. Once identified, the core of the CSSI pipeline, based on an iterative statistical hypothesis-testing procedure [16], is used to cluster all nucleolus voxels, followed by a max-voting strategy to separate them into distinct nucleolar volumes. Finally, the nucleolar and nuclear volumes are utilized to extract multiplexed quantitative subcellular measurements from the cell’s 3D RI tomogram.

A detailed description of the BPC-based CSSI nucleolus segmentation algorithm is provided in the Supplementary Information (Section S1 and Table S1) and illustrated for an experimental HeLa cell (Fig. S1).

### Assessment of the nucleoli segmentation

To assess the proposed nucleoli segmentation method within 3D RI tomograms of flowing/suspended cells recorded by the HTFC system, HeLa cells were used as an experimental model. The HTFC setup and numerical processing described in the Methods section were applied to reconstruct the 3D RI tomograms of 224 flowing/rotating HeLa cells [11]. For each cell, hundreds of QPMs (Fig. S1(a,b)) were extracted from the holographic video, the unknown viewing/rolling angles were retrieved, and tomographic reconstruction was performed (Fig. S1(c)). Stain-free nucleoli were then identified using the CSSI algorithm operating on the 3D BPC space (Fig. S1). Representative tomograms during cell flow and rotation are shown in Fig. 3(a,b), illustrating a typical HTFC acquisition. In Fig. 3(a), segmentation performed using four RI thresholds isolates volumes that do not correspond to specific intracellular structures, confirming the need for advanced segmentation in HTFC. In contrast, Fig. 3(b) shows that the proposed method successfully extracts both nucleus and nucleoli. Examples of tomograms with different nucleoli counts per cell are provided in Fig. 3(c).

**Fig. 3.**
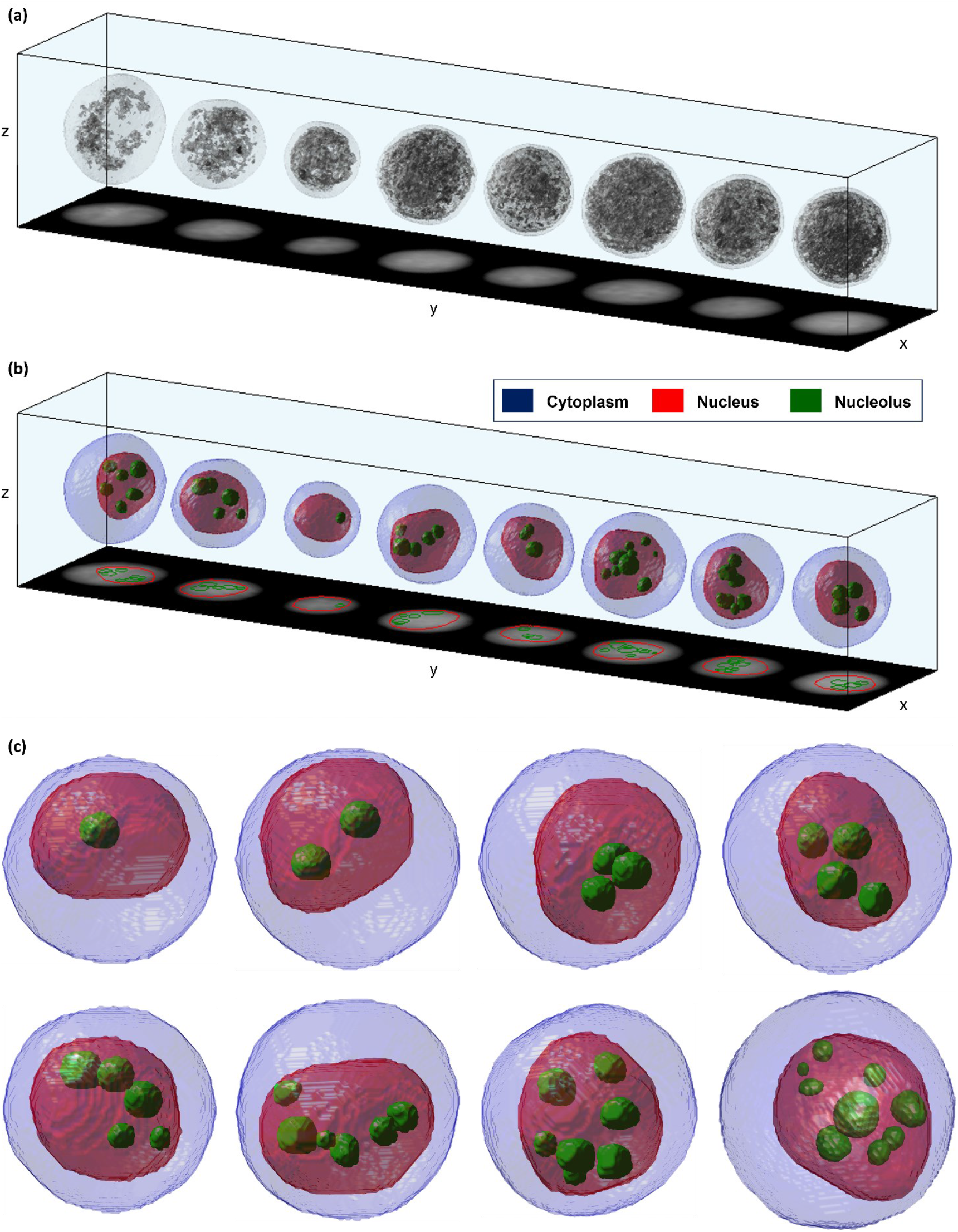
Examples of 3D HTFC tomograms of HeLa cells (Supplementary Movie S1). **(a,b)** Tomograms of the same single cells flowing and rotating inside the microfluidic channel. In (a), isolevels obtained by using four RI thresholds do not correspond to specific intracellular structures. In (b), the nucleus (red) and the nucleoli (green) have been segmented through the CSSI algorithms. For each tomogram, the corresponding 2D QPM along the z-axis is reported at the bottom of the channel. **(c)** Atlas of cells with different numbers of nucleoli (green) within the nucleus (red).

To evaluate these results, a direct comparison between stain-free HTFC and in-flow 3D FM on the same cells would be ideal. However, such a multimodal system is not currently existing as it is cumbersome to realize. Therefore, validation was performed indirectly by comparing quantitative nucleolar measurements with corresponding FM data acquired on the same cell line across different modalities. Specifically, three assessments were carried out (see Section S2 and Table S2 in Supplementary Information for details): (i) comparison with previously reported static 2D FM measurements [29]; (ii) comparison with data obtained here using a commercial in-flow 2D FM cytofluorimeter; and (iii) comparison with data acquired here using a static 3D FM confocal microscope.

In 224 HeLa cells, HTFC yielded 3.3 ± 1.6 nucleoli per cell (mode = 3), consistent with static 2D FM measurements reported in the literature (3.5 ± 1.7 with mode = 3), as shown in Fig. 4(a) [29]. Instead, using a commercial in-flow 2D FM cytofluorimeter, 325 cells were analyzed with Draq-5 (nucleus) and anti-Fibrillarin (nucleoli) staining (Fig. 4(b,c)), showing 3.3 ± 1.8 nucleoli per cell (Fig. 4(d)). Both in-flow 2D FM and HTFC data exhibited the same mode of 3 nucleoli per cell, and a Student’s t-test confirmed no significant difference between the two distributions (p-value = 0.88). In fact, the Student’s t-test is a statistical method used to compare two datasets [37], and the resulting p-value indicates the probability that the observed difference arises by chance. Hence, higher p-values suggest a greater likelihood that the two datasets come from the same population.

**Fig. 4.**
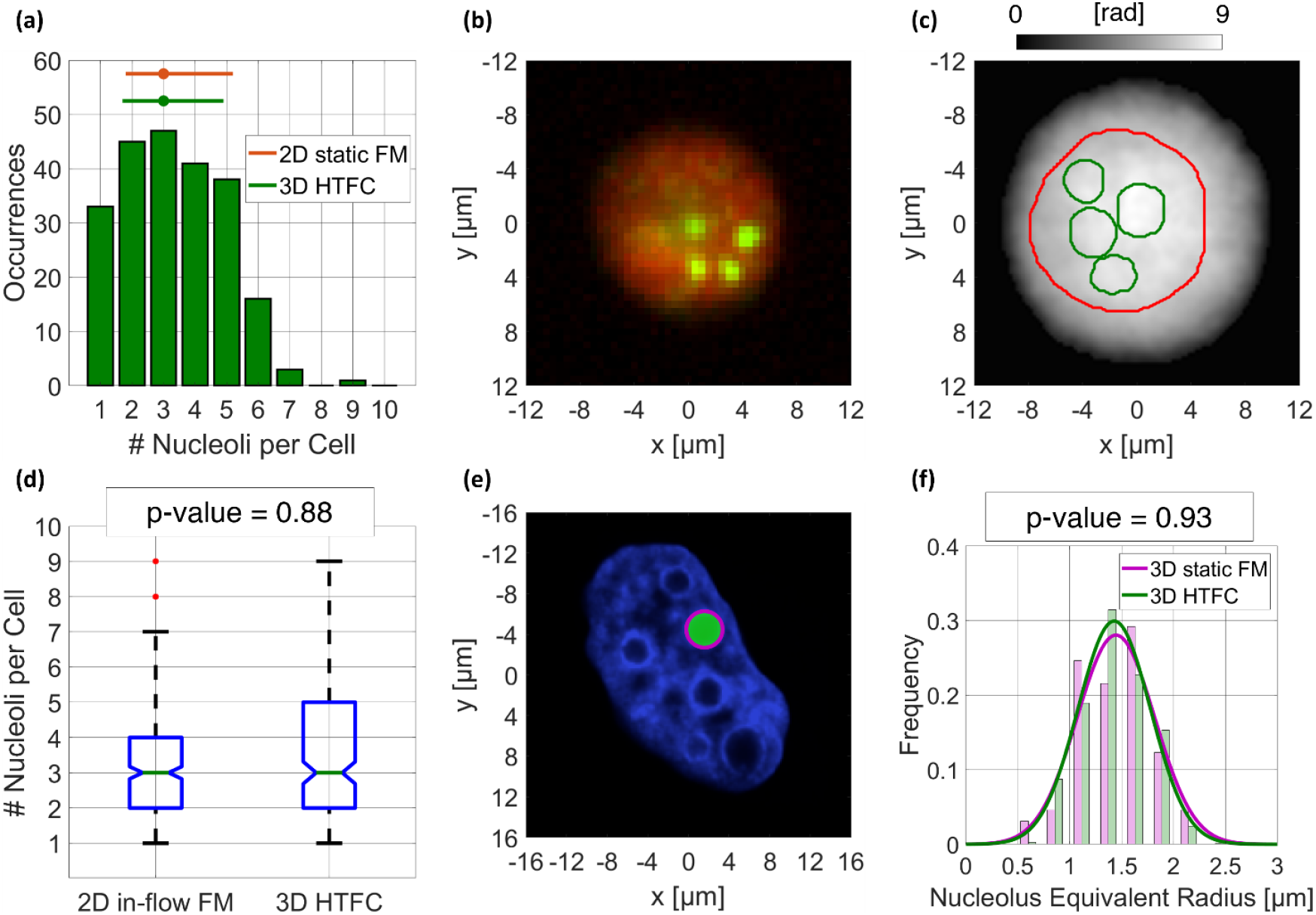
Assessment of the nucleoli segmentation in HTFC by comparison with standard FM techniques within a dataset of HeLa cells. **(a)** Histogram of the number of nucleoli per cell measured in 2D static FM (50,087 cells) [29] and in 3D HTFC (224 cells), with the corresponding statistics at the top (bar is the mean ± standard deviation and dot is the mode value). **(b)** Cell imaged by 2D in-flow FM cytofluorimetry with the nucleus and nucleoli marked in red and green, respectively. **(c)** QPM with overlapped the contours of the nucleus (red) and nucleoli (green) obtained by reprojecting the HTFC tomogram in Fig. S1(n) segmented by the BPC-based CSSI algorithm. **(d)** Boxplot of the number of nucleoli per cell measured in 2D in-flow FM cytofluorimetry (325 cells) and in 3D HTFC (224 cells). In each box, the central mark indicates the median, while the top and bottom edges represent the 75th and 25th percentiles, respectively. The notch around the median provides a visual indication of the 95% confidence interval. Whiskers extend to the most extreme points not considered outliers. **(e)** Central slice of a cell imaged by 3D static FM confocal microscopy with the nucleus marked in blue and with overlapped an example of elliptic approximation for measuring the nucleolus size (green). **(f)** Histogram of the nucleolus size measured in 3D static FM confocal microscopy (65 nucleoli in 20 cells) and in 3D HTFC (744 nucleoli in 224 cells), with overlapped the corresponding gaussian fitting.

For 3D validation, high-content static 3D FM confocal microscopy was performed, staining the nucleus with Hoechst dye. From 65 nucleoli in 20 HeLa cells, the equivalent nucleolar radius was 1.434 ± 0.348 µm, in excellent agreement with 3D HTFC measurements from 744 nucleoli in 224 HeLa cells (1.430 ± 0.319 µm), with no statistically significant difference (p-value = 0.93), as shown in Fig. 4(f).

These results collectively demonstrate that the BPC-based CSSI nucleolus segmentation achieves quantitative accuracy comparable to state-of-the-art FM while maintaining fully label-free operation.

### Multiplexed measurements of single-cell intracellular/intranuclear biophysical properties

The main advantage of HT over FM confocal microscopy lies in its ability to provide quantitative biophysical features derived from the 3D RI distribution at the single-cell level [30]. Moreover, unlike standard 3D static FM confocal microscopy and conventional HT, HTFC operates on suspended cells rather than adherent ones, enabling the analysis of the most reliable intracellular and intranuclear 3D organization without the distortions introduced by surface adhesion. The flow cytometry module additionally facilitates the acquisition of large datasets for the statistically significant measurement of entire cellular populations while preserving single-cell variability. In this framework, the implementation of CSSI algorithms enables, for the first time, the multiplexed study of statistical distributions of biophysical features extracted from 3D HTFC tomograms of flowing/suspended cells after retrieving specificity for nucleus and nucleoli, as sketched in Fig. 2(a). To this end, we analyzed a dataset of 224 HeLa cells comprising 744 nucleoli.

The histogram of the mean RI for each segmented nucleolus is shown in Fig. 5(a). Variations in organelle RI have been linked to several pathologies [38], and differences in RI across intracellular compartments can offer new insight into cell biology. For this reason, Fig. 5(b–d) reports the histograms of the ratios between the mean RI contrast of nucleolus and nucleoplasm, nucleolus and cytoplasm, and nucleoplasm and cytoplasm, respectively. In particular, here the RI contrast is defined as the difference between the mean RI of a compartment and that of the surrounding medium (*n*_0_ = 1.334), the nucleoplasm corresponds to the nuclear volume not occupied by nucleoli, and the cytoplasm corresponds to the cell volume not occupied by the nucleus. As expected, the nucleolus exhibits a higher RI than the surrounding nucleoplasm [30,38], which in turn shows a higher RI than the cytoplasm in this cell line.

**Fig. 5.**
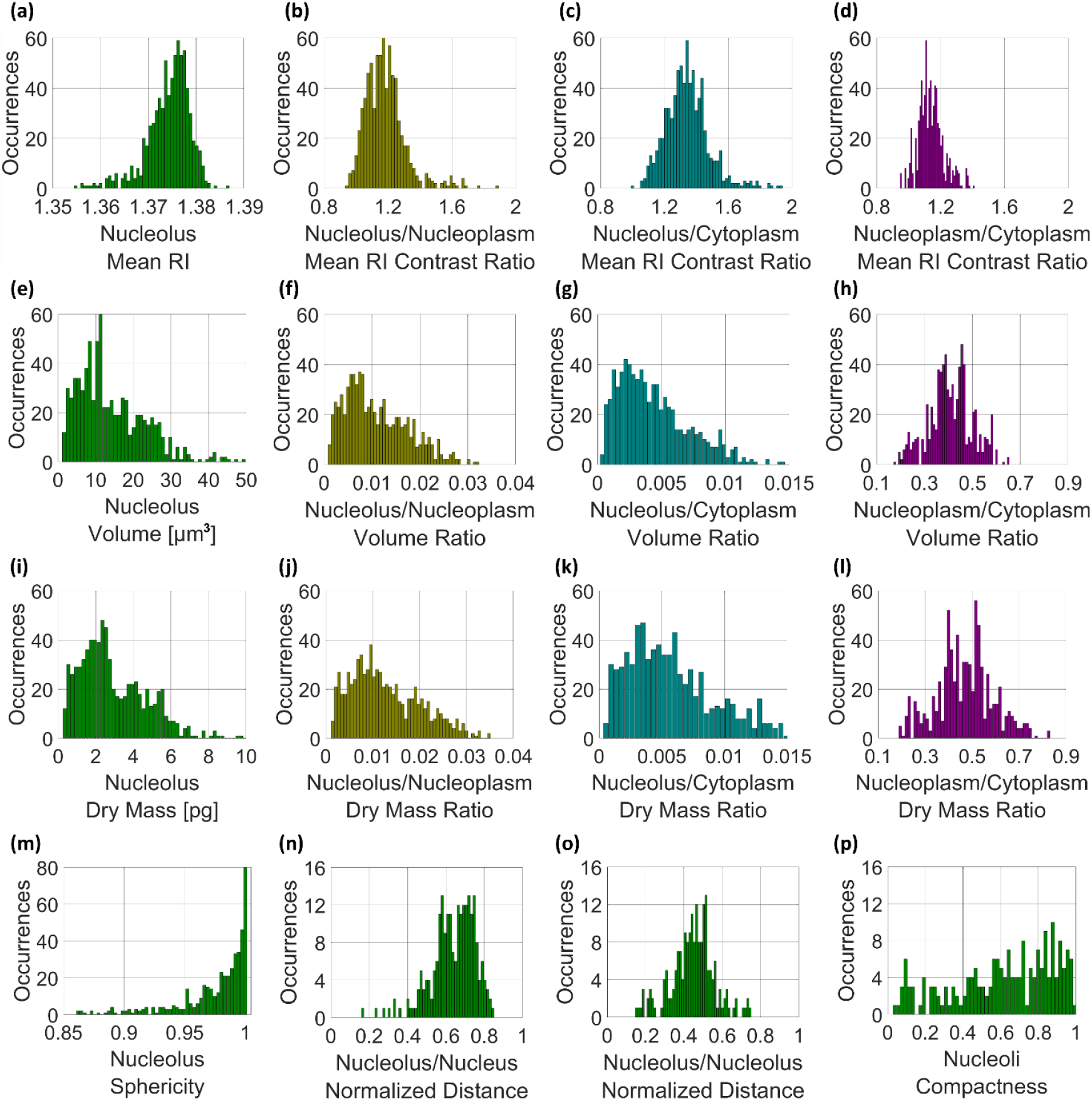
Histograms of multiplexed biophysical features computed from a dataset of 224 HeLa cells and corresponding 744 nucleoli after retrieving the intracellular/intranuclear specificity within the 3D HTFC tomograms. **(a)** Mean RI of the nucleolus. **(b)** Mean RI contrast ratio between the nucleolus and the nucleoplasm. **(c)** Mean RI contrast ratio between the nucleolus and the cytoplasm. **(d)** Mean RI contrast ratio between the nucleoplasm and the cytoplasm. **(e)** Volume of the nucleolus. **(f)** Volume ratio between the nucleolus and the nucleoplasm. **(g)** Volume ratio between the nucleolus and the cytoplasm. **(h)** Volume ratio between the nucleoplasm and the cytoplasm. **(i)** Dry mass of the nucleolus. **(j)** Dry mass ratio between the nucleolus and the nucleoplasm. **(k)** Dry mass ratio between the nucleolus and the cytoplasm. **(l)** Dry mass ratio between the nucleoplasm and the cytoplasm. **(m)** Sphericity of the nucleolus. **(n)** Average distance between the nucleoli of each cell and the nucleus centroid, normalized to the nucleus equivalent radius. **(o)** Average distance among all the nucleoli of each cell, normalized to the nucleus equivalent diameter. **(p)** Compactness of the nucleoli inside the nucleus.

HTFC tomograms also provide access to intracellular volumetric features. In fact, nucleolar size (Fig. 5(e)), in addition to nucleolar count (Fig. 4(a)), are recognized hallmarks of cancer when increased [39,40]. Thus, monitoring them is important for assessing therapeutic strategies targeting nucleolar function [25,39]. Conversely, reduced nucleolar number and size correlate with increased longevity [25,41]. In Fig. 5(f,g), nucleolar volume is normalized to nucleoplasm and cytoplasm volumes, respectively, while Fig. 5(h) reports the nucleoplasm-to-cytoplasm volume ratio, a well-established cancer biomarker [42-44].

By combining the average RI with the compartment volume, the dry mass can be computed as one of the hallmark biophysical quantities accessible through HT [5], since it reflects the non-aqueous mass of an organelle. The histogram of nucleolar dry mass is shown in Fig. 5(i), while Fig. 5(j,k) reports its normalization to the dry mass of the nucleoplasm and cytoplasm, respectively. The nucleoplasm-to-cytoplasm dry mass ratio is shown in Fig. 5(l).

Beyond nucleolar count and size, nucleolar morphology is also a recognized cancer biomarker. While nucleoli are typically roundish in healthy cells, irregular shapes have been reported in malignant ones [45-47]. Recording suspended cells with HTFC enables accurate measurement of nucleolar sphericity (Fig. 5(m)), defined as the ratio between the surface area of a sphere with the same nucleolar volume and the actual nucleolar surface area (0 for non-spherical objects, 1 for perfect spheres).

Recovering multiplexed specificity also allows quantifying the true 3D spatial organization of organelles. Nucleolar positioning within the nucleus carries functional significance and remains an open question in cell biology [48-50]. Here, we propose several descriptors of intranuclear architecture. For each cell, we compute the average nucleolus–to–nucleus-centroid distance normalized to the nucleus equivalent radius (Fig. 5(n)), showing that nucleoli tend to reside closer to the nuclear membrane than to its centroid. Fig. 5(o) shows the normalized nucleolus–nucleolus distance, defined as the mean inter-nucleolar spacing normalized to the nucleus equivalent diameter. In this HeLa dataset, the average distance between two nucleoli corresponds to ∼45% of the nucleus equivalent diameter. Finally, Fig. 5(p) reports nucleoli compactness, describing how nucleoli are spatially spread within the nucleus. For each cell, it is computed as the ratio between the volume of nucleoli convex hull, defined as the intersection between the nucleus and the smallest sphere enclosing all nucleoli, and the nucleus volume. On average, the nucleoli convex hull occupies 62% of the nuclear volume.

Average and standard deviation values for all features in Fig. 5 are summarized in Table S3.

## Conclusion

Despite the strong appeal of label-free microscopy techniques (including QPI), the lack of intrinsic intracellular specificity has long represented the main bottleneck for its widespread adoption. In continuity with our recent efforts to extend HTFC specificity to multiple organelles (nucleus, lysosomes, lipid droplets, vacuoles), in this work we introduced a novel metric space based on a 3D pseudo-RI tomogram that enhances the contrast of small intracellular structures. Applying CSSI to this pseudo-tomogram enables the extraction of complex intranuclear organelles such as nucleoli that remain unresolved in conventional HTFC 3D RI reconstructions. This capability stems from the proposed BPC transformation, which compensates for optical aberrations caused by the quasi-spherical morphology of suspended cells in flow, effectively converting standard RI maps into a representation where sub-nuclear features become distinctly detectable.

The method was validated on a large dataset of HeLa cells using multiple fluorescence-based techniques (static 2D FM, in-flow 2D FM, and static 3D FM), showing strong quantitative consistency and no statistically significant difference in terms of nucleoli count and size. These results demonstrate that BPC-based CSSI provides fluorescence-comparable specificity and quantitative reliability in a fully 3D label-free context.

Beyond nucleolar segmentation, this framework supports multiplexed intranuclear and intracellular characterization, enabling simultaneous quantification of cytoplasm, nucleoplasm, and nucleoli in live suspended cells. This represents a meaningful advancement for HTFC, expanding its capabilities to multi-organelle analysis.

Overall, transforming the conventional 3D RI space into a BPC representation unlocks new quantitative possibilities for HTFC, narrowing the long-standing specificity gap with FM. By combining volumetric label-free imaging with statistical throughput, the proposed strategy provides a robust foundation for future single-cell diagnostic and prognostic applications, particularly in cancer and degenerative disease research.

### Online Methods

#### Sample preparation

HeLa wild type cells were maintained in Dulbecco’s modified Eagle’s medium (DMEM) containing 10% fetal bovine serum (FBS), 200mM L-Glutamine, 100 U/ml penicillin and 100 μg/ml streptomycin.

For 2D in-flow FM cytofluorimetry, 7 million of HeLa cells were collected by a wash with phosphate-buffered saline (PBS) 1× and incubation with a 0.05% trypsin–EDTA solution (Sigma). After centrifugation for 5 min at 125xg, cells were resuspended in PBS 1× and exposed to the anti-Fibrillarin primary antibody (ab4566) at a dilution rate of 1:500 for 90 min. Then, the staining with AlexaFluor 488 goat anti-mouse (A28175, Invitrogen) secondary antibody was performed for 45 min. At the end, cells were incubated with 25 µM DRAQ5 fluorescent probe (#62254, Thermo Scientific) for 5 min at room temperature.

For 3D FM confocal microscopy, HeLa cells were washed with PBS 1×, detached with trypsin-EDTA and then plated in 96 wells at the concentration of 8000 cells/well for 24 h. The day after, cells were fixed with PFA 4%, washed with PBS 1×, and then stained with Hoechst for 10 min at room temperature. For HTFC, Hela cells underwent a similar preparation process. Cell counting was performed with Trypan blue, and the cell suspension was adjusted to a concentration of 300,000 cells. During the HTFC acquisition, cells were maintained in a suspended and live state.

#### 2D in-flow FM cytofluorimetry

Amnis^®^ ImageStream^®^X Mark II flow cytometer (Luminex) was used to acquire single-cell images at 40× magnification. The acquired raw image file (.rif) contained among 1000 events (10–30 events per second). The analysis of single cells fluorescence intensity was performed by using IDEAS software (version 6.2.64.0). To consider only single cells, a dot plot showing area versus aspect ratio was created. Then, we generated a morphology mask that defined nucleus stained by DRAQ5 and the feature “spot count” was used to estimate the number of nucleoli in the HeLa nucleus based on Fibrillarin intensity.

#### 3D FM confocal microscopy

HeLa cells were plated on 96-well plates (PerkinElmer, ViewPlate cod. 6055302) and then acquired using high-content imaging instrumentation, specifically the Opera Phenix system (PerkinElmer). Images were acquired with 63× magnification and with 5 different focus positions (Z-stack, distance between planes = 0.5 µm). At the end of the acquisition, the morphological parameters of nucleoli were measured with the Harmony analysis software (PerkinElmer).

#### HTFC experiments

To perform HTFC experiments, a digital holographic microscope based on a Mach Zehnder interferometer in off-axis configuration was implemented, as sketched in Fig. 6(a) [11]. Cells are injected into the microfluidic channel (Microfluidic ChipShop 10000107 – L_z_ = 200 μm, L_x_ = 1000 μm, L_y_ = 58.5 mm) by an automated syringe pump (Cetoni NEMESYS 290 N) which allows controlling flow with a quasi-uniform speed (∼50 nl/s). By exploiting the hydrodynamic forces of the laminar flow, cells move along the y-axis and mainly rotate along the x-axis. Light wave emitted by the laser (Laser Quantum Torus, 532 nm) is split into an object beam and a reference beam by a polarizing beam splitter (PBS). Furthermore, two half-wave plates (HWPs) are placed in front of and behind the PBS for balancing the intensity ratio of the two split contributions. Scattering produced by the roto-translating cells illuminated by the object beam is collected by a microscope objective (MO1, Zeiss Plan-Apochromat 40× NA=1.3 Oil immersion) and sent to a tube lens (TL1, focal length 150 mm). At the same time, the reference beam follows a free path passing through a beam expander shaped by a second microscope objective (MO2, NewPort 20× NA=0.40) and a second tube lens (TL2, focal length 250 mm). Finally, the two beams are recombined by a beam splitter (BS) with a non-null angle to obtain off-axis configuration. The resulting interference patterns propagate up to a CMOS camera (Genie Nano-CXP Cameras 5120×5120 pixels, pixel size equal to 4.5 μm), which records the holographic video sequence at 30 fps over a field of view (FoV) of 640 μm × 640 μm.

**Fig. 6.**
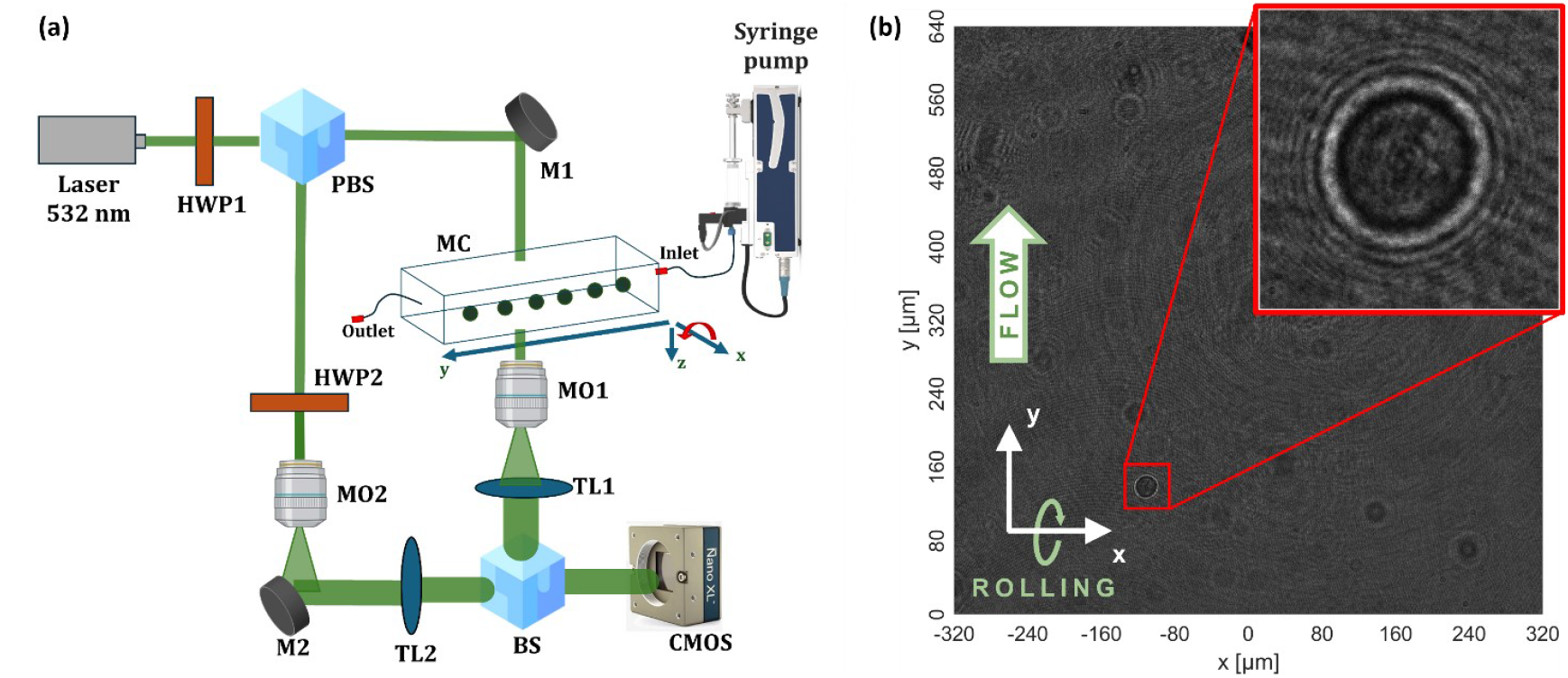
HTFC system for recording holographic video sequences. **(a)** Sketch of the implemented Mach-Zehnder setup. HWP: Half Wave Plate; PBS: Polarizing beam splitter; M: mirror; MO: microscope objective; TL: Tube Lens; MC: microfluidic channel; BS: beam splitter; CMOS: camera. **(b)** Example of captured 5120×5120 digital hologram, from which the 384×384 holographic ROIs are cropped (red inset).

An example of digital hologram is displayed in Fig. 6(b). From each captured 5120×5120 FoV, multiple regions of interest (ROIs) sized at 384×384 pixels are isolated, as that highlighted in Fig. 6(b). Each holographic ROI is then converted into the corresponding QPM by means of numerical processing [11]. Specifically, the holographic ROI is initially demodulated utilizing a Fourier-domain bandpass filter. The demodulated hologram is then numerically propagated along the optical z-axis using the Angular Spectrum formula to determine the cell focal distance by minimizing a contrast-based metric. In fact, in digital holography, usually cells are captured out of focus as they can be numerically refocused post-experiment. After refocusing the complex field of the cell, its argument is extracted. Starting from the retrieved phase values, any remaining aberrations are compensated for by subtracting a reference hologram recorded without any sample in the microfluidic channel. Subsequently, denoising and unwrapping algorithms are applied to obtain the final QPM (Fig. S1(a)). Hence, a stack of hundreds of QPMs at different viewing angles is computed for each roto-translating cell. After estimating the unknown viewing angles from the y-positions, the 3D RI tomogram is reconstructed by using the Filtered Back Projection algorithm [11].

## Data and Code availability

Data and code will be made available from the Corresponding Author upon reasonable request.

## Acknowledgements

This work was partially supported by project PRIN 2022 PNRR - Flow-cytometry ImaGing by Holographic tomography for predicting TUMor control in Oncology patients treated with Radiotheraphy (FIGHT-TUMOR), Prot. P2022ATE2J – funded by the Italian Ministry of University & Research in the framework of Next Generation EU.

This work was partially supported by project “Advanced Cancer diagnosis: Combining label-freE fLow cytometry techniques EmpoweRed by ArTificial IntElligence (ACCELERATE)”, CUP B53C22006670001, within *Bandi a cascata Progetto ANTHEM (Code PNC_0000003) Spoke 2* – funded by the Italian Ministry of University & Research in the framework of Next Generation EU. This work was partially supported by project “FIGHT-LSDs” – Flow-cytometry ImaGing by Holographic Tomography to advance study, diagnosis and treatment of Lysosomal Storage Diseases – funded by *Accordo per la Coesione della Regione Campania. Fondo di Rotazione ex L. 183/1987* (project n. 48).

Dr. Martina Mugnano contributed to the manuscript when she was affiliated to the Department of Chemical, Materials and Production Engineering, DICMaPI, University of Naples “Federico II”, Piazzale Tecchio 80, 80125 Naples, Italy. Currently, she is affiliated to the CNR-IGB, Institute of Genetic and Biophysics “Adriano Buzzati Traverso”, Via Pietro Castellino 111, 80131 Naples, Italy.

