## Supplementary Information for "Label-Free Nucleoli Measurement by 3D Holo-Tomographic Flow Cytometry Using Biolens Phase Compensation"

### S1. BPC-based CSSI nucleolus segmentation algorithm

In this section, the several steps of the CSSI nucleolus segmentation algorithm based on the BPC method (Fig. 2) are described in detail and illustrated in Fig. S1 by using an experimental HeLa cell recorded through the HTFC system. As reported in the Methods, hundreds of QPMs (Fig. S1(a,b)) are obtained for each cell from the recorded holographic video sequence and, after retrieving the corresponding unknown viewing/rolling angles, the tomographic reconstruction is performed (Fig. S1(c)). However, as illustrated in the central slice of the HeLa cell nucleus (Fig. S1(c)), the intranuclear RI contrast is insufficient to automatically identify the nucleolus reference voxel clouds with high confidence due to biolens phase aberrations introduced by the quasi-spherical cell. In fact, the quasi-spherical morphology of a suspended cell generates a quasi-paraboloid wavefront in the 2D QPM [9,26]. This characteristic is evident in the pseudo-3D visualization shown in Fig. S1(b), where the curvature of the wavefront hides fine details of intranuclear features such as nucleoli. Starting from the recorded 2D QPM, the BPC method is implemented to obtain the corresponding 2D pseudo-QPM, as shown in Fig. S1(d,e). Comparison between the pseudo-3D rendering of the QPM and pseudo-QPM in Fig. S1(b,e), respectively, clearly reveals the improvement in intracellular contrast (see black arrow). This enhancement is further confirmed in Fig. S1(g), where the normalized QPM and pseudo-QPM intensity profiles corresponding to the lines in Fig. S1(a,d), respectively, are directly compared, highlighting the improved contrast achieved thanks to the BPC. Thus, biolens phase aberrations are compensated in each 2D QPM through the BPC method.

1. The BPC method is implemented for all the 2D QPMs related to the same cell. In particular,
   a. considering all the $L_x \times L_y$ QPMs related to the same cell, the maximum height $H$ and the maximum width $W$ are computed;
   b. a $W \times H$ region of interest (ROI) is cropped around the centre of each QPM;

c. the third order polynomial fitting is computed for each $W \times H$ QPM ROI;
d. the $W \times H$ quasi-paraboloid is subtracted to each $W \times H$ QPM ROI, in order to obtain the corresponding $W \times H$ pseudo-QPM ROI;
e. each pseudo-QPM ROI is reported to the original $L_x \times L_y$ size by zero-padding, thus obtaining the corresponding pseudo-QPM;
f. the sequence of pseudo-QPMs and corresponding viewing/rolling angles is given in input to the tomographic algorithm;
g. within the reconstructed tomogram, the sole nucleus is isolated;
h. within the isolated nucleus, the background values are set as the RI of the surrounding medium in the HTFC experiments (i.e., $n_0$), while $n_0 + 0.01 - M_N$ is added to all the cell values, where $M_N$ is the minimum nucleus value, thus obtaining the nucleus 3D pseudo-RI tomogram, which central slice is shown in Fig. S1(f).

By comparing the central slice of the original 3D RI tomogram (Fig. S1(c)) and the corresponding 3D pseudo-RI tomogram mapped in the BPC space (Fig. S1(f)), the visual localization of nucleolar compartments within the nucleus is much more evident. This property is also underlined in Fig. S1(h), where the RI and pseudo-RI profiles related to the lines in Fig. S1(c,f), respectively, are compared after normalization between 0 and 1. Hence, this intranuclear contrast enhancement permits to implement an automatic searching of their reference sets, which are groups of voxels belonging to the organelles to be segmented, used as a reference for the subsequent statistical comparisons. At this aim, the known features about nucleoli can be exploited. Indeed, it is well known that nucleoli have a quasi-spherical shape [35] with the highest RI values within the nucleus [30]. Moreover, this latter property has been enhanced in the BPC space.

2. $S$ searching elements are created as two concentric spheres with radii $R_{int}$ and $R_{ext}$, where
   a. the internal sphere has a homogeneous pseudo-RI equal to the average value among the nucleus pseudo-RIs greater than their $T$ percentile;
   b. the external sphere has a homogeneous pseudo-RI equal to the average value among the nucleus pseudo-RIs less than their $T$ percentile;
   c. the background has a homogeneous RI equal to $n_0$;
   d. the internal spheres have $S$ possible radii $R_{int} \in [R_1; R_2]\ \mu m$ with uniform step $\Delta R = (R_2 - R_1)/(S - 1)$;
   e. the external spheres have radii $R_{ext} = R_{int} + 2\Delta R$.
3. For each of the $S$ searching elements,
   a. it is used as template to compute the 3D normalized cross-correlation (NCC) with respect to the nucleus 3D pseudo-RI tomogram [36];
   b. a binary cross-correlation tomogram is created by selecting the real values of the 3D NCC greater than its maximum real value multiplied by $T$;
4. the logical $or$ is performed between the $S$ binary cross-correlation tomograms, thus obtaining a preliminary nucleoli reference sets tomogram;

The 3D NCC is exploited to find the local regions within the nucleus 3D pseudo-RI tomogram that match well with the searching elements, which have been simulated as concentric spheres in order to emulate the typical scenario of a nucleolus, that is a sphere having high pseudo-RI values (i.e., the nucleolus) immersed in a lower pseudo-RI distribution (i.e., the nucleus), in turn immersed within the background medium. In this way, the 3D NCC provides the highest values in those points where the local 3D pseudo-RI distribution is more similar to the nucleolus template (see the central slice of a

3D NCC in Fig. S1(i)). Furthermore, since no exact a priori information about the nucleolus size is available, the nucleolus templates are simulated with different sizes (i.e., the $S$ searching elements). For this reason, having enhanced the intranuclear contrast through the quasi-paraboloid subtraction is a crucial step for carrying out this analysis in a more reliable way. However, the 3D NCC could give in output also some localization error.

5. The high RI values of the nucleoli are exploited again for filtering the outlier reference sets, i.e., within the preliminary nucleoli reference sets tomogram, the reference sets having an average pseudo-RI less than the $T$ percentile of the nucleus 3D pseudo-RI distribution are removed, thus obtaining the filtered nucleoli reference sets tomogram.
6. The watershed algorithm [S1] is employed to separate the reference sets that are too close each other, thus avoiding that they are considered as one single reference set in the following CSSI analysis.

After identifying the $N$ reference sets of the nucleoli (see Fig. S1(j)), the core of the CSSI algorithm can be implemented.

7. For each reference set, steps 1-5 reported in the Supplementary Information of Ref. [16] are implemented, which can be briefly summarized as follows.
   a. A searching volume (see Fig. S1(k)) is defined as the intersection between the nucleus volume and a sphere centred in the reference set with a radius equal to half the equivalent radius of the surrounding nucleus (i.e., the radius of a sphere having the same nucleus volume);
   b. the 3D pseudo-RI tomogram of the analyzed nucleus is centered in its $L_x \times L_y \times L_z$ array and then divided into distinct cubes, each of which has an edge measuring $\varepsilon$ pixels (step 1 in Ref. [16]);
   c. the pseudo-RI values of each cube are compared to the pseudo-RI values of the reference set through a non-parametric hypothesis statistical test, i.e. the Wilcoxon-Mann-Whitney (WMW) test [S2,S3] (steps 2-3 in Ref. [16]);
   d. cubes having the highest p-values are clustered into a preliminary nucleolus set (steps 2-3 in Ref. [16]);
   e. the preliminary nucleolus set is filtered and refined, in order to obtain a partial nucleolus set, which is made of nucleolus sub-cubes with an edge measuring $\varepsilon/2$ pixels (steps 4-5 in Ref. [16]);
   f. steps a-e are repeated $K$ times, thus obtaining $K$ partial nucleolus sets, that are slightly different from each other because, to make more robust the statistical clustering, in some of the several WMW tests, a random selection of the reference set from the original one is performed.

Hence, at this stage, $K$ partial nucleolus sets are associated with each of the $N$ starting reference sets. Of course, it could happen that the same nucleolus sub-cube has been assigned to the partial nucleolus sets of different reference sets, above all when the reference sets are close each other (see Fig. S1(l)).

8. To delete this ambiguity, a max-voting strategy can be exploited thanks to the existence of $K$ partial nucleolus sets for each reference set. In fact, each clustered nucleolus sub-cube is assigned to the reference set for which it occurred more times. In this way, for each reference set, the $K$ partial nucleolus sets converge to one single final nucleolus set (see Fig. S1(m)).
9. For each final nucleolus set,
   a. a spatial and statistical filtering of the nucleolus sub-cubes is performed (step 4 in Ref. [16]);

b. all the possible pairs of remaining sub-cubes are linked through a line segment (step 6 in Ref. [16]);
c. a morphological closing is performed to smooth the corners and fill the holes of the resulting 3D polygonal (steps 7 in Ref. [16]), thus obtaining the nucleoli segmentation (see Fig. S1(n)).

By comparing Fig. S1(n) with Fig. S1(j), it can be noted that the number of reference sets is 5, while the number of segmented nucleoli is 4 because some false positives detected during the identification of the reference steps are automatically deleted by the following clustering steps, since no statistical similarity is found within the nucleus able to converge to a well-defined nucleolus. All the parameters involved in the described BPC-based CSSI nucleolus segmentation algorithm are reported in Table S1.

### S2. Comparison with multiple fluorescence references

*Comparison with 2D static FM measurements*

From HTFC measurements, we observed $3.3 \pm 1.6$ nucleoli per cell, as shown in the histogram in Fig. 4(a). Of note, this measurement appears in line with the one reported in the literature. In particular, a robust study about the number of nucleoli in several cell lines, carried out using a standard 2D FM imaging system, reported $3.5 \pm 1.7$ nucleoli per cell in HeLa cells [29]. By comparing the statistics about the 2D static FM and 3D HTFC techniques at the top of Fig. 4(a), the segmentation obtained through the proposed BPC-based CSSI method fits well with the measurements reported in literature [29]. Above all, both the measurements exhibit the same mode value, i.e. 3 nucleoli per cell (Fig. 4(a)).

*Comparison with 2D in-flow FM cytofluorimetry measurements*

We also employed a commercial in-flow 2D cytofluorimeter (i.e., Amnis® ImageStream®X Mark II) to record the FM images of 325 HeLa cells in which the Draq-5 and anti-Fibrillarin antibody were used to mark the nucleus and the nucleoli, respectively. An example of 2D FM image is displayed in Fig. 4(b), in which 4 nucleoli can be clearly observed in green with respect to the surrounding nucleus (red). To visually compare the 2D in-flow FM image with a segmented 3D HTFC tomogram, the latter must be first converted to a 2D image. To do this, we exploited as example the 3D RI tomogram segmented in Fig. S1(n). In particular, we considered a stain-free QPM taken from the overall stack used for the tomographic reconstruction, in which the intracellular compartments cannot be recognized due to the lack of exogeneous markers (Fig. 4(c)). Then, after reconstructing the 3D RI tomogram and implementing the CSSI algorithms for the nucleus and nucleoli identification, we reprojected the segmented tomogram along the same viewing direction as the selected QPM in order to mark the contours of the nucleus (red) and its 4 nucleoli (green). Hence, the 2D QPM (Fig. 4(c)) can be computationally segmented and visually compared with the 2D in-flow FM image (Fig. 4(b)). It is important to note that, by combining HTFC to the CSSI algorithms, not only the intracellular and intranuclear specificity can be retrieved, but also a biophysical characterization of the cellular compartments can be obtained. Indeed, the quantitative properties related to propagation of the light through the sample can be measured from the 2D QPM in Fig. 4(c) [5], which is instead unfeasible within the 2D FM image in Fig. 4(b).

In addition to the simple visual assessment, a statistical comparison between the 2D in-flow FM and 3D HTFC experiments can be performed because of the large number of imaged cells through the flow cytometry modality. In particular, in the boxplots of Fig. 4(d), we reported the number of nucleoli measured with the 2D in-flow FM images (i.e., $3.3 \pm 1.8$) and the 3D HTFC images (i.e., $3.3 \pm 1.6$) of 325 and 224 HeLa cells, respectively. Also in this case, the two histograms exhibit the same mode, i.e. 3 nucleoli per cell. Moreover, by performing a Student's t-test between the two distributions [37], a very high p-value was obtained (i.e., 0.88), thus meaning that the two measurements obtained by 2D in-flow FM and 3D HTFC have been extracted from the same distribution with high probability.

*Comparison with 3D static FM confocal microscopy measurements*

Finally, in order to consider the inherently 3D nature of the HTFC technique, we also performed experiments through a high-content static 3D FM confocal microscope by staining the nucleus with the Hoechst dye. An example of 3D static FM image is displayed in Fig. 4(e), in which the nucleoli can be clearly seen as roundish organelles (black) within the marked nucleus (blue). From the 3D static FM experiments, $3.3 \pm 0.9$ nucleoli per cell were measured, again with a mode of 3 nucleoli per cell. However, we exploited the 3D measurements to perform an additional statistical assessment of the nucleoli segmentation within the 3D HTFC data based on the nucleolar sizes. In fact, starting from the segmented 3D RI tomograms, we measured the volume of each nucleolus, from which we computed the equivalent radius, i.e. the radius of a sphere having the same volume as the segmented nucleolus. As regards the 3D static FM images, for each nucleolus we selected the central slice and we measured the area of its elliptic approximation (Fig. 4(e)). In fact, in 3D static FM, cells are adherent over a substrate and not suspended as in 3D HTFC, as can be inferred from the elongated shape of the nucleus in Fig. 4(e). Nevertheless, nucleoli preserve their roundish shape due to their very smaller size with respect to the overall cell volume (Fig. 4(e)). For this reason, the elliptic fitting can be considered a good approximation of the nucleolar volume. In this way, for each 3D static FM nucleolus, we computed the equivalent radius as the radius of a circle having the same area as its elliptic approximation. Following this strategy, we measured an equivalent radius of $1.434 \pm 0.348$ μm from the 65 nucleoli of 20 HeLa cells imaged by 3D static FM and $1.430 \pm 0.319$ μm from the 744 nucleoli of 224 HeLa cells imaged by 3D HTFC (Fig. 4(f)). The good agreement between the two histograms in Fig. 4(f) about the nucleolar size is confirmed by the high p-value (i.e., 0.93) obtained through the Student's t-test [37].

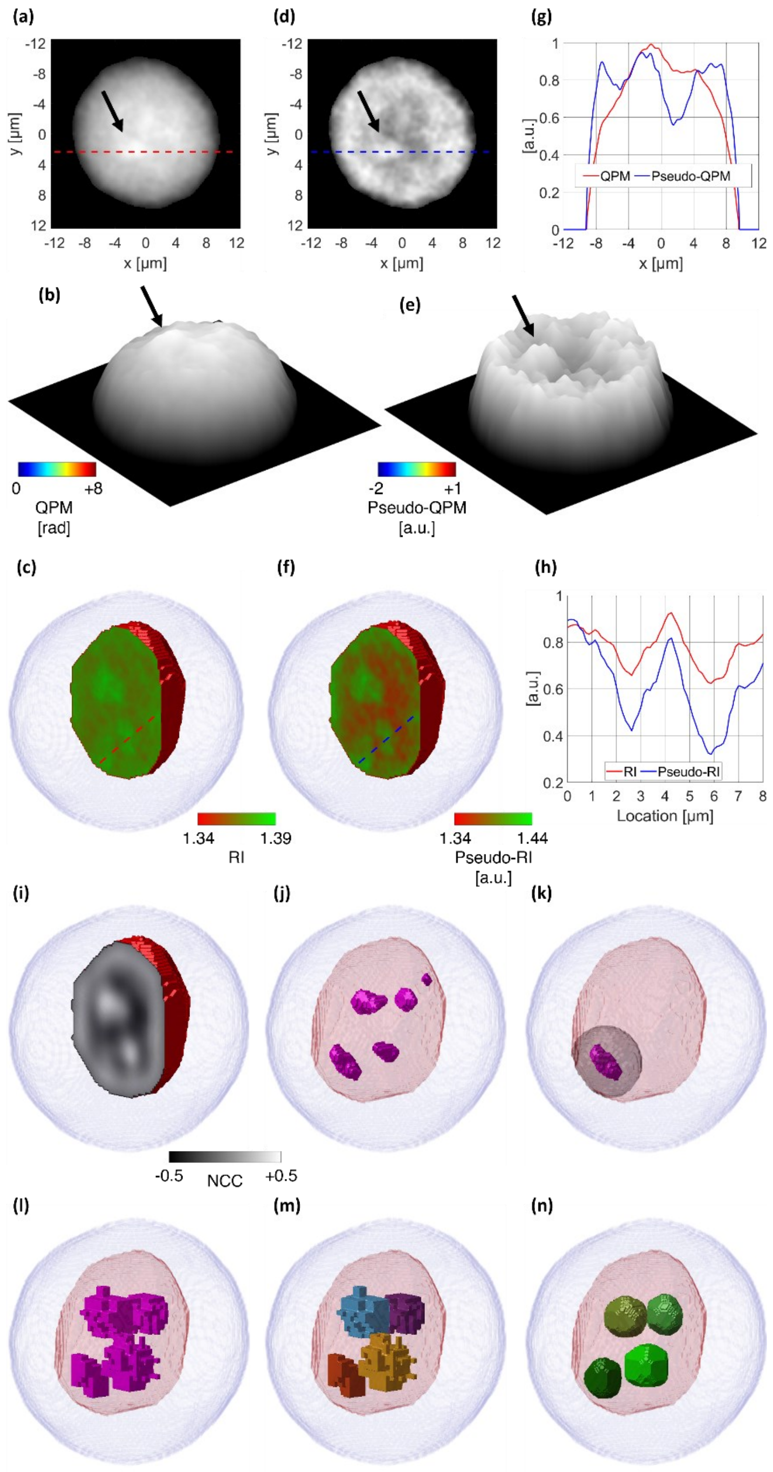


**Fig. S1. Segmentation of nucleoli within a HeLa cell. (a)** QPM taken from the overall QPM sequence of the flowing/rolling cell. **(b)** Pseudo-3D visualization of the QPM in (a). **(c)** Central slice of the nucleus 3D RI tomogram within the cell shell. **(d)** Pseudo-QPM frame obtained by removing the quasi-paraboloid pattern from the QPM in (a) through the BPC method. **(e)** Pseudo-3D visualization of the pseudo-QPM in (d). In (a,b,d,e), the black arrows highlight the contrast enhancement about the nucleolus. **(f)** Central slice of the nucleus 3D pseudo-RI tomogram within the cell shell, visualized in the new 3D metric space obtained by the BPC method. **(g)** Normalized QPM-profile (red) and pseudo-QPM profile (blue) taken from the lines in (a,d), respectively. **(h)** Normalized RI profile (red) and pseudo-RI profile (blue) taken from the lines in (c,f), respectively. **(i)** Central slice of the 3D NCC obtained by setting $R_{int} = 2R_1$. **(j)** Reference sets (magenta) for the nucleoli segmentation within the nucleus (red) and cell (light blue) shells. **(k)** Searching volume (grey) related to the bottom-left reference set (violet) in (h). **(l)** Overall sub-cubes (violet) belonging to the $K$ partial nucleolus sets associated with all the $N$ starting reference sets. **(m)** Sub-cubes divided for each starting reference set (orange, yellow, cyan, and violet) by means of max-voting. **(n)** Nucleoli (different green colors) segmented by the BPC-based CSSI algorithm within the nucleus (red) and cell (light blue) shells.

| **Table S1.** Parameters used in the proposed BPC-based CSSI algorithm to segment the stain-free nucleoli from the 3D RI tomograms of flowing cells recorded by HTFC. | | | |
|---|---|---|---|
| $L_x = L_y = L_z = 200\ pixels$ | $n_0 = 1.334$ | $T = 0.75$ | $R_1 = 0.5\ \mu m$ |
| $R_1 = 2\ \mu m$ | $S = 7$ | $\varepsilon = 8\ pixels$ | $K = 16$ |

**Table S2.** Comparison between the measurements of nucleoli segmented in HTFC by the BPC-based CSSI algorithm with respect to the corresponding ones related to nucleoli marked in standard FM techniques within a dataset of HeLa cells.

| | # Cells | # Nucleoli | # Nucleoli per cell | Nucleolus size [µm] |
|---|---|---|---|---|
| 2D static FM [29] | 50087 | 175304 | $3.5 \pm 1.7$ | - |
| 2D in-flow FM cytofluorimetry | 325 | 1072 | $3.3 \pm 1.8$ | - |
| 3D static FM confocal microscopy | 20 | 65 | $3.3 \pm 0.9$ | $1.434 \pm 0.348$ |
| 3D HTFC | 224 | 744 | $3.3 \pm 1.6$ | $1.430 \pm 0.319$ |

| **Table S3.** Multiplexed biophysical features computed from a dataset of 224 HeLa cells and corresponding 744 nucleoli after retrieving the CSSI-based intracellular/intranuclear specificity within the 3D HTFC tomograms. | |
|---|---|
| Nucleolus Mean RI | $1.374 \pm 0.005$ |
| Nucleoplasm Mean RI | $1.368 \pm 0.005$ |
| Cytoplasm Mean RI | $1.364 \pm 0.004$ |
| Nucleolus/Nucleoplasm Mean RI Contrast Ratio | $1.182 \pm 0.126$ |
| Nucleolus/Cytoplasm Mean RI Contrast Ratio | $1.343 \pm 0.135$ |
| Nucleoplasm/Cytoplasm Mean RI Contrast Ratio | $1.140 \pm 0.079$ |
| Nucleolus Volume [$\mu m^3$] | $901.4 \pm 567.6$ |
| Nucleoplasm Volume [$\mu m^3$] | $8370.6 \pm 1817.2$ |
| Cytoplasm Volume [$\mu m^3$] | $2118.9 \pm 5322.4$ |
| Nucleolus/Nucleoplasm Volume Ratio | $0.011 \pm 0.007$ |
| Nucleolus/Cytoplasm Volume Ratio | $0.004 \pm 0.003$ |
| Nucleoplasm/Cytoplasm Volume Ratio | $0.409 \pm 0.090$ |
| Nucleolus Dry Mass [pg] | $2.9 \pm 1.8$ |
| Nucleoplasm Dry Mass [pg] | $233.9 \pm 51.5$ |
| Cytoplasm Dry Mass [pg] | $522.1 \pm 136.8$ |
| Nucleolus/Nucleoplasm Dry Mass Ratio | $0.013 \pm 0.007$ |
| Nucleolus/Cytoplasm Dry Mass Ratio | $0.006 \pm 0.004$ |
| Nucleoplasm/Cytoplasm Dry Mass Ratio | $0.469 \pm 0.120$ |
| Nucleolus Sphericity | $0.984 \pm 0.028$ |
| Nucleolus/Nucleus Normalized Distance | $0.639 \pm 0.115$ |
| Nucleolus/Nucleolus Normalized Distance | $0.448 \pm 0.113$ |
| Nucleoli Compactness | $0.616 \pm 0.267$ |
